# Kinetic and Structural Analysis of Ribonucleotide Extension by DNA Polymerase η

**DOI:** 10.1101/2022.09.12.507597

**Authors:** Caleb Chang, Christie Lee Luo, Sarah Eleraky, Aaron Lin, Grace Zhou, Yang Gao

## Abstract

DNA polymerases catalyze DNA synthesis with high fidelity, which is essential for all life. Extensive kinetic and structural efforts have been executed in exploring mechanisms of DNA polymerases, surrounding their kinetic pathway, catalytic mechanisms, and factors that dictate polymerase fidelity. Recent time-resolved crystallography studies on DNA polymerase η (Pol η) and β have revealed essential transient events during DNA synthesis reaction, such as mechanisms of primer deprotonation, separated roles of the three metal ions, and conformational changes that disfavor incorporation of the incorrect substrate. DNA-embedded ribonucleotides (rN) are the most common lesion on DNA and a major threat to genome integrity. While kinetics of rN incorporation has been explored and structural studies have revealed that DNA polymerases have a steric gate that destabilizes rNTP binding, mechanism of extension upon rN addition remains poorly characterized. Using steady-state kinetics, static and time-resolved X-ray crystallography with Pol η as a model system, we showed that the extra hydroxyl group on the primer terminus does not reduce the catalytic efficiency of Pol η. However, rN ended primers alter the dynamics of the polymerase active site as well as the catalysis and fidelity of DNA synthesis. During rN extension, Pol η fidelity drops significantly across different sequence context. Systematic structural studies suggest that the rN at the primer end improved primer alignment and reduced barriers in C2’-endo to C3’-endo sugar conformation change. Our work provides important insights for rN extension and implicates a possible mechanism for rN removal.

## Introduction

DNA polymerases are given the essential task of high-fidelity replication of genomic DNA. Seventeen DNA polymerases from seven enzymatic families have been identified in human cells responsible for DNA replication, DNA damage bypass, and DNA repair (1,2). Although distinct in their sequences and structures, DNA polymerases contain similar active sites, follow similar kinetic pathways, and employ similar catalytic mechanisms (1). Multiple metal ions are required for promoting DNA synthesis (**Fig. 1a**) (3–6): the B-site metal ion (Me^2+^_B_) is associated with the triphosphate motif of the incoming deoxynucleotide triphosphate (dNTP) and stabilizes its binding; the A-site metal ion (Me^2+^_A_) lies between the primer end and the incoming dNTP and aligns the primer 3’-OH with the substrate α-phosphate to promote primer 3’-OH deprotonation and nucleophilic attack; following the binding of dNTP and the Me^2+^_A_ and Me^2+^_B_, the C-site metal ion (Me^2+^_C_) binds between the α- and β-phosphates on the other side of the active site and drives α-β-phosphate bond breakage. Accompanied with product formation, the primer terminus sugar pucker changes from a C2’-endo to C3’-endo conformation to avoid steric clashes (3). Upon product formation, the metal ions and pyrophosphate dissociate from the polymerase active site and the newly synthesized primer end translocates out of the nucleotide insertion site for the next round of dNTP incorporation.

**Fig. 1:**
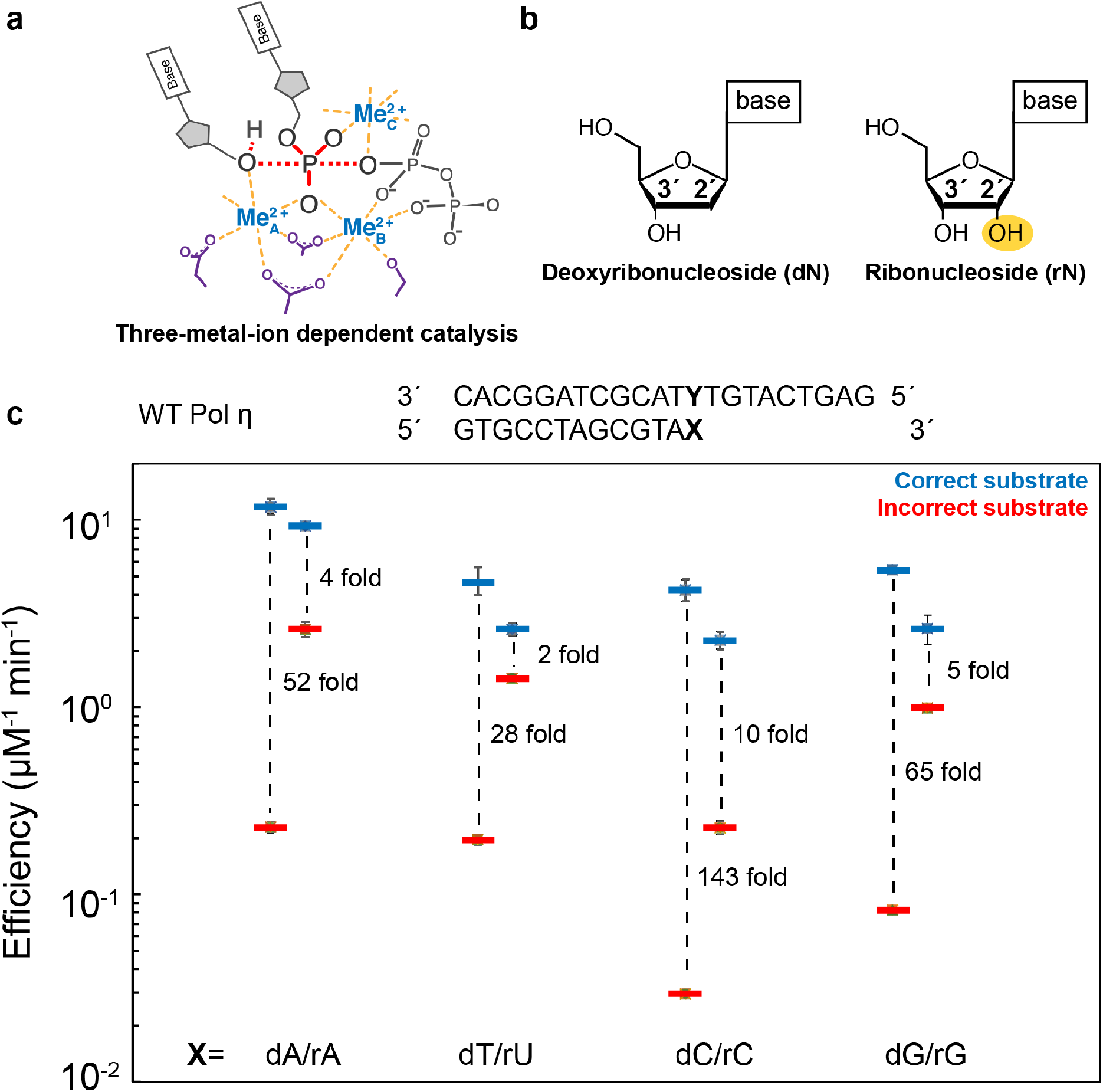
Ribonucleotide extension. **a**, Scheme of the three-metal-ion dependent enzyme catalysis and transition state stabilization. Divalent metal ions optimally bind to their ligands with bond lengths of 2.0-2.2 Å at 90 °. **b**, Chemical representation of deoxyribonucleoside (dN) and ribonucleoside (rN). Hydrogens on the sugar moieties have been removed for clarity. **c**, WT Pol η DNA polymerase fidelity during dN or rN extension in the presence of Mg^2+^ based on steady-state kinetics. The bars represent the mean of duplicate measurements for the catalytic efficiencies (k_cat_/K_M_) for incorporation of dATP (blue) and dGTP (red) opposite dT for WT Pol η. The errors bars represent the standard deviation for the measurements. The distance between the respective catalytic efficiencies is a measure of discrimination.

Genomic DNA constantly faces endogenous and environmental assaults. Although there exist proficient repair pathways, polymerases inevitably face DNA lesions while traveling along DNA. Such lesions may create bulky obstacles or alter base-pairing, pi-stacking, and the DNA backbone, promoting polymerase stalling and replication stress (7). Eukaryotic cells contain repair and translesion polymerases that can bypass various lesions (2). Different translesion mechanisms are used to bypass different lesions with varying chemical properties (2). DNA-embedded ribonucleotides (rN) are the most common lesions and pose major threats to genome integrity (8,9). A vast majority of rNs are incorporated during DNA replication by DNA polymerases δ and ε (10,11). In addition, rNs are incorporated by Pol α as primers during Okazaki fragment formation (12). During NHEJ or base excision repair, rN incorporation by X-family polymerases facilitates downstream ligation (13,14). Having a OH group on the 2’ carbon (**Fig. 1b**), DNA embedded rNs can lead to nicks on the phosphate backbone and the accumulation of nonsynonymous mutations (11,15). These rNs, if left incorporated on the nuclear or mitochondria template strand, can lead to error-prone impairment and stalling of the replication and transcription machinery (16–19). Because of their lethal nature, the cell has evolved two specific pathways to remove DNA-embedded rNs. The first involves ribonuclease H2 (RNaseH2), which nicks at the rN site for error-free ribonucleotide excision repair (RER) (20,21). The second involves topoisomerase I, which can lead to small DNA deletions (22–25). In addition, mismatch repair (MMR) has been implicated as an alternative pathway in rN removal (26), but the mechanism of rN recognition is unclear.

The kinetic and structural basis of rN incorporation by DNA polymerases is well understood (19). In A-family (27,28), B-family (29,30), and Y-family polymerases (31–33), a steric gate formed by tryptophan and phenylalanine residues sterically clash with the hydroxyl group on the 2’ carbon of ribo-nucleotide triphosphates (rNTP) and prevent rNTP binding. X-family polymerases such as DNA polymerase μ, β, and λ contain only a peptide backbone in place of the steric gate and thus exhibit much lower sugar pucker discrimination (34). Notably, DNA polymerase μ prefers to incorporate dNTP over rNTP by only 2-fold (35). During DNA replication, if a rN does get incorporated, DNA synthesis extending the rN can still occur as the rN contains a functional 3’-OH group. Recently, it was revealed that the catalytic rate of DNA polymerase ε drops 3300 fold during rN extension compared to deoxy-nucleotide (dN) extension (36), hinting the involvement of translesion polymerases in rN extension. Kinetic analysis of Pol β revealed insignificant changes in catalytic efficiency (K_cat_/K_M_) between dN and rN extension (37). Structural studies of DNA polymerase λ showed minimal differences when the 3’-end of primer was a dN versus a rN during correct nucleotide incorporation (38). Despite such efforts, whether and how the 2’-OH at the primer end affects the structure and dynamics of the active site as well as polymerase catalysis and fidelity are not fully explored.

DNA polymerase η (Pol η) is a Y-family translesion polymerase responsible for bypassing cyclobutane pyrimidine dimers (39,40). Besides, Pol η has been implicated in translesion DNA synthesis against a variety of lesions (2). People with mutations on the Pol η gene develop a predisposition for skin cancer and xeroderma pigmentosum (41,42). In addition, Pol η widely participates during the synthesis of the lagging strand (43) and has the ability to incorporate rNs and exhibits reverse transcriptase activity on RNA:DNA duplexes (44). Pol η has been used as a model system for investigating mechanisms of polymerase catalysis. Kinetic studies have revealed that Pol η follows a similar kinetic pathway to other polymerases (45). The reaction process of Pol η was captured at atomic resolution with recent time-resolved crystallography, which has revealed numerous insights into the fundamental mechanisms of DNA synthesis, including roles of metal ions, factors that control fidelity, and deprotonation mechanism of the primer 3’-OH (3–5,46–48). In addition, structural snapshots of various lesion bypass processes by Pol η were captured for illustrating mechanisms of translesion synthesis (40,49–51). Thus, we sought to use the Pol η system to investigate the consequences of rN extension at atomic resolution.

Here, we present biochemical and structural studies of Pol η extending rNs. Steadystate kinetic data on correct and incorrect single-rN extension suggest that Pol η can extend rNs with high efficiency. Interestingly, the rN primer end significantly lowers replication fidelity. Corresponding crystal structures of Pol η complexed with both Mg^2+^ and Mn^2+^ and single rN-primed DNA substrate suggest that having a rN at the primer terminus stabilizes it in a productive conformation for nucleophilic attack. Furthermore, we compare the misincorporation process extending from ribose uridine (rU) and deoxyribose thymine (dT) primed DNA substrate with time-resolved crystallography. The results further confirm that the decreased Pol η fidelity during rN extension is due to the stabilization of the active aligned conformation during nucleophilic attack. Our biochemical and structural studies of Pol η provide novel insights for rN extension and suggest implications for rN removal.

## Results

### Kinetics and fidelity of rN extension by Pol η

During the DNA synthesis reaction, the primer 3’-OH is aligned with the α-phosphate for nucleophilic attack. It was revealed that primer 3’-OH alignment promoted by Me^2+^_A_ is the key step in substrate discrimination (5,52). Interestingly, Gregory and coworkers captured the rN primer terminus in the aligned conformation in the absence of the Me^2+^_A_ (48), suggesting the additional 2’-OH group may alter conformation of the sugar ring and possibly facilitate nucleophilic attack. We thus hypothesized that a rN at the 3’-primer end may promote primer 3’-OH alignment and lower DNA synthesis fidelity. Steady-state kinetic assays of Pol η with native and single-rN primed DNA substrate were conducted to detect for changes in fidelity during rN extension. The kinetic assays indicated that correct nucleotide incorporation efficiency during rN and dN extension was roughly similar (**Fig. 1c**). However, misincorporation efficiency from a rN primer was enhanced and the extension fidelity was reduced by over 10-folds. Previous studies suggested that the fidelity of Pol η is sequence dependent and decreases when extending from deoxyribose adenine (dA) and deoxyribose thymine (dT), also known as the TA motif (53). We thus also examined DNA substrates with different terminal nucleotides. Similar as reported, Pol η catalyzed DNA synthesis with different fidelity on substrates with different primer termini, with deoxyribose cytosine (dC) as the highest and dA/dT as the lowest. For all sequence context, the fidelity dropped over 10 fold with a rN primer end. Notably for rU extension, correct incorporation was only twice more efficient than misincorporation during rN extension.

### Fidelity of S113A Pol η

Previous crystal structures of Pol η showed that the primer is coordinated in the down conformation by serine 113 in the active site (**Fig. 2a**). This widely conserved serine among Y-family polymerases is important for primer alignment (48). We investigated whether a S113A Pol η mutant would perturb primer alignment and enhance polymerase fidelity. With a dN primer end, the S113A mutant catalyzed DNA synthesis with 77-fold fidelity, higher than that of the wild-type (WT), confirming the role of S113. From the discrimination plot in **figure 2b**, S113A Pol η fidelity dropped by 6-fold when a rN-ended primer was used. A rN at the primer end stimulated correct extension by 2-fold, but incorrect extension by 17-fold. This was consistent with previous studies, where ribose adenine (rA) at the primer termini overwrites the effect of S113 to help in aligning the primer for nucleophilic attack. The results further confirmed the significant role of primer alignment in fidelity control and the conformation selection of the rN primer end.

**Fig. 2:**
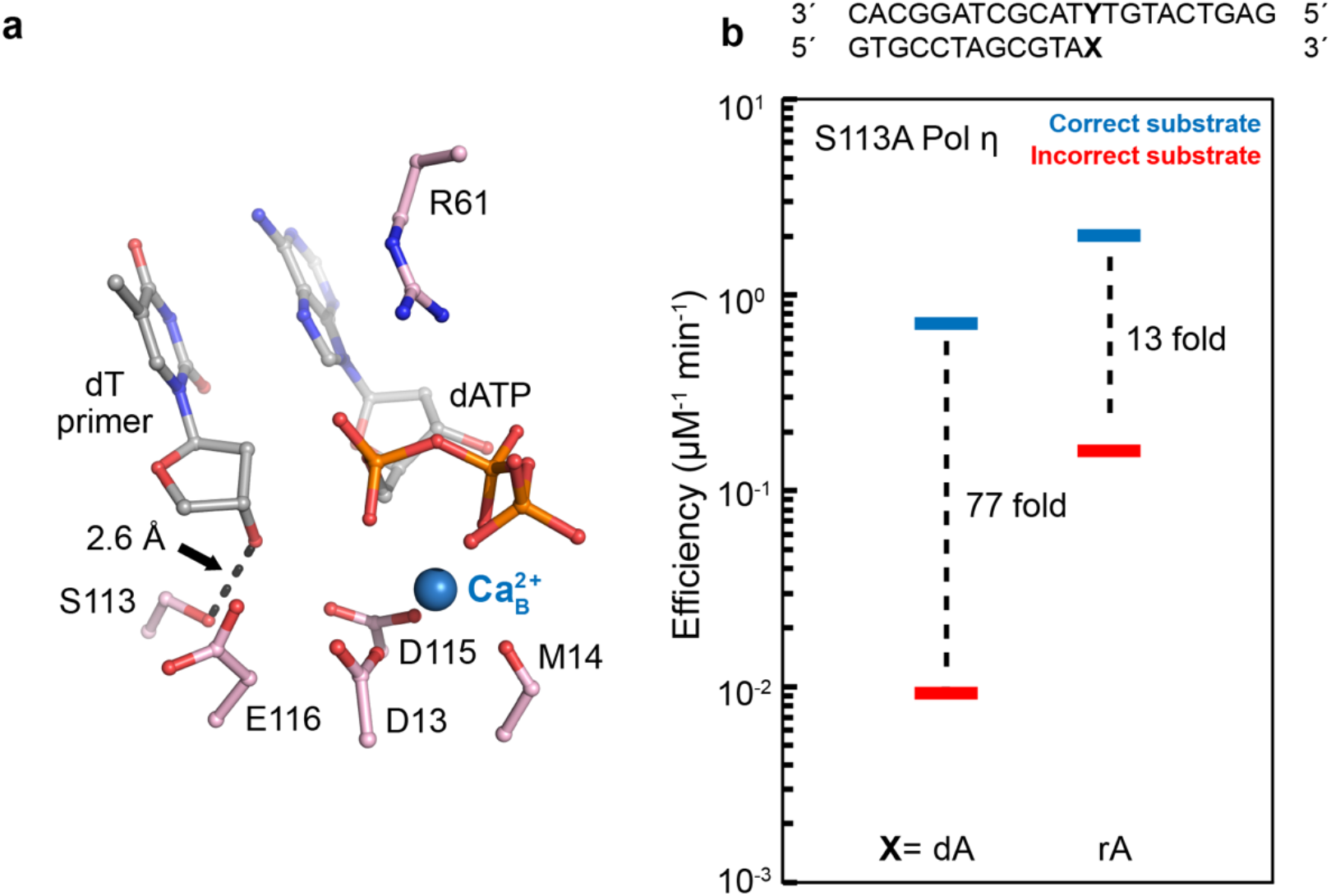
Mechanisms of S113A polymerase η catalysis and misincorporation. **a**, Pol η ground state with the primer terminus close to S113 (PDB ID **4ECQ**). **b**, S113A Pol η DNA polymerase fidelity during dN or rN extension in the presence of Mg^2+^ based on steady-state kinetics. The bars represent the mean of duplicate measurements for the catalytic efficiencies (k_cat_/K_M_) for incorporation of dATP (blue) and dGTP (red) opposite dT.

### Binary complex of Pol η with dN and rN ended primer

To test how the rN primer end would affect primer alignment, we first obtained crystals of Pol η in the absence of incoming nucleotide. We captured the binary state of Pol η with dT at 1.75 Å resolution and rU at 2.1 Å resolution (**Fig. 3**). Both structures are similar to the previous ternary complex (RMSD 0.2 and 0.3, respectively) and binary complex (RMSD 0.3 and 0.4, respectively), confirming that there are no significant conformational changes during incoming nucleotide binding (54). The dT and rU structures are almost identical. In addition, the sugar ring conformations are the same. The sugar ring for both primers in the binary complex is in a C2’ endo geometry. The 3’-OH group exists 6.8 Å and 6.3 Å away from the S113 within the dT and rU structures, respectively. These binary structures suggested that the primer 3’-ends with either dN or rN are not aligned in the absence of the incoming dNTP and Me^2+^.

**Fig. 3:**
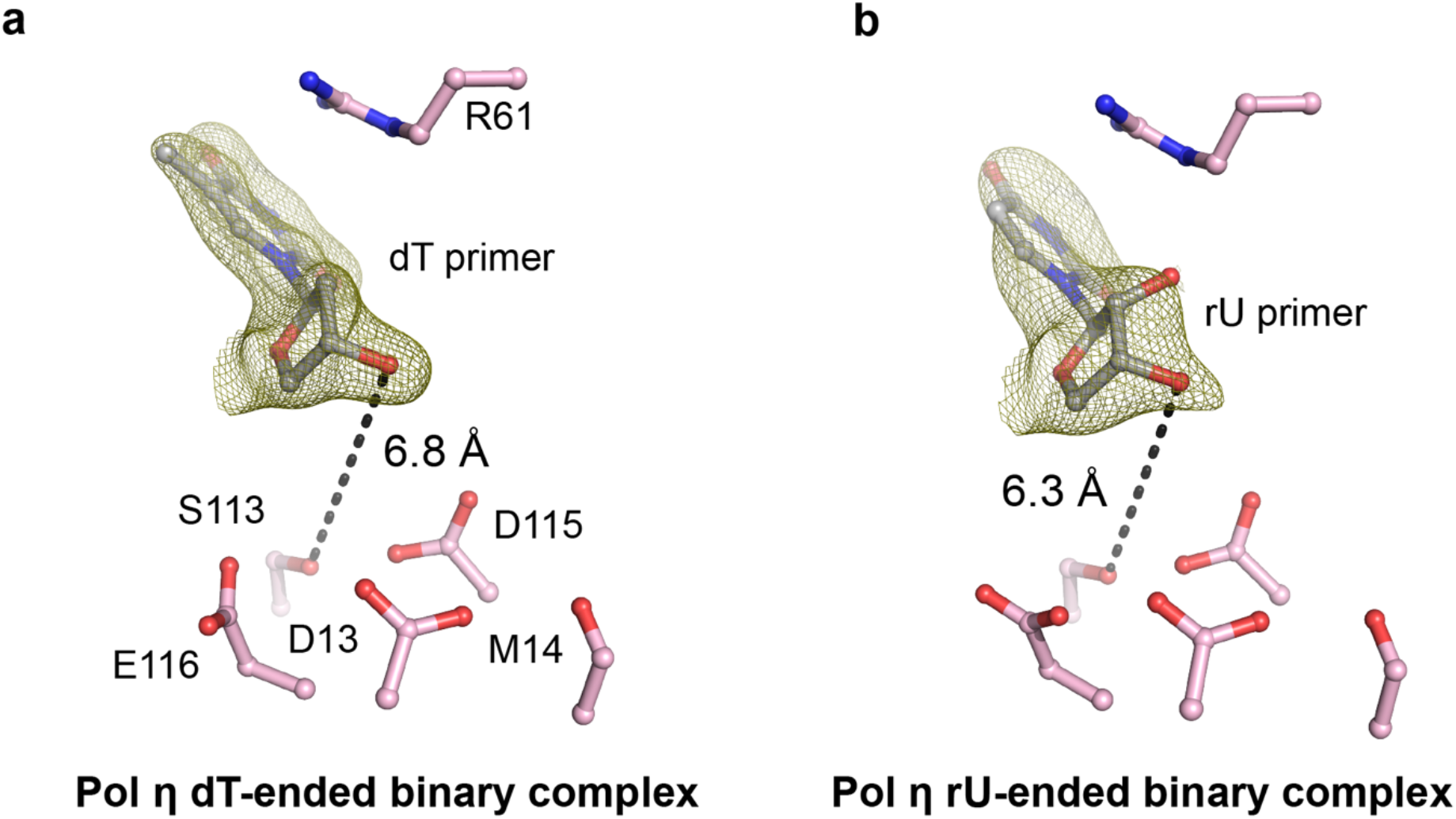
Pol η:DNA binary structure. Binary structure of Pol η complexed with dT-primed in **a** or rU-primed DNA in **b**. The F_o_-F_c_ omit map for the primer (green) was contoured at 3 σ (σ values represent r.m.s. density values).

### *Structures of* Pol η *misincorporation complex with rN at the primer terminus*

We further investigated how a rN primer end affects primer alignment by capturing the ternary misincorporation complex. To prevent catalysis, we prepared ternary Pol η structures with rN-ended primers complexed with 2’-deoxyguanosine-5’-[(α,β)-imido]triphosphate (dGMPNPP) against a dT template. We determined structures of Pol η with rA, rU, rC, ribose guanine (rG) at the primer terminus. In the rA structure, the primer is 100% aligned even with the incorrect substrate, in contrast to only 25% in the dA structure (**Fig. 4a, b**) (5). It is interesting to note that for the rA primed structure, the sugar pucker of the rA base was already in a C3’endo conformation, as opposed to that of the dA structure, which was in a C2’ endo conformation (**Fig. 4a and 5**). Similarly, the rU structure was with 100% of the rU primer in a C3’endo conformation and in the aligned down conformation (**Fig. 4c, d**). In contrast, the primer termini in the rC and rG structures existed in the up non-productive conformation, similar to the structures with dC and deoxyribose guanine (dG) (**Fig. 6a,b and Supplementary Fig. 1**) (53).

**Fig. 4:**
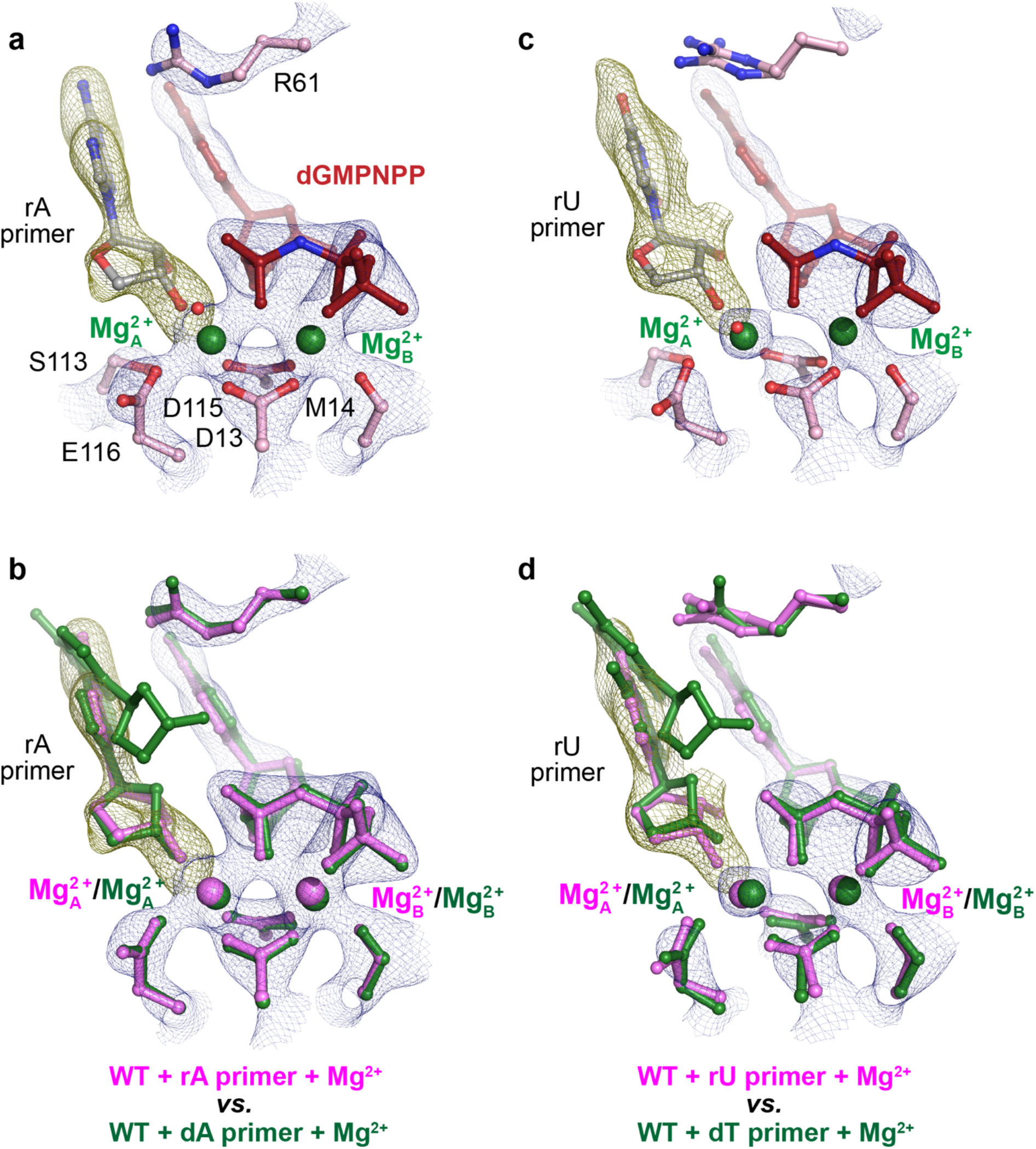
Structural comparison between reactant states of incorrect nucleotide incorporation during dN or rN extension. **a, c**, Structure of rA extension (**a**) and rU extension (**c)**WT Pol η mismatch reactant state complexed with Mg^2+^. **b**, **d,** Structural overlay of dN (green) (PDB ID **4J9M** and **4J9K**) or rN (pink) extension structures with dGMPNPP:dT base-pair indicating differences in 3’-OH alignment. All electron density maps apply to the molecule colored in pink. The 2F_o_-F_c_ map for everything including Me^2+^_A_ and Me^2+^_B_, dGMPNPP, and catalytic residues and S113 (blue) was contoured at 2 σ. The F_o_-F_c_ omit map for the primer terminus (green) was contoured at 3.5 σ in **a, b** and 2.7 σ in **c, d**.

**Fig. 5:**
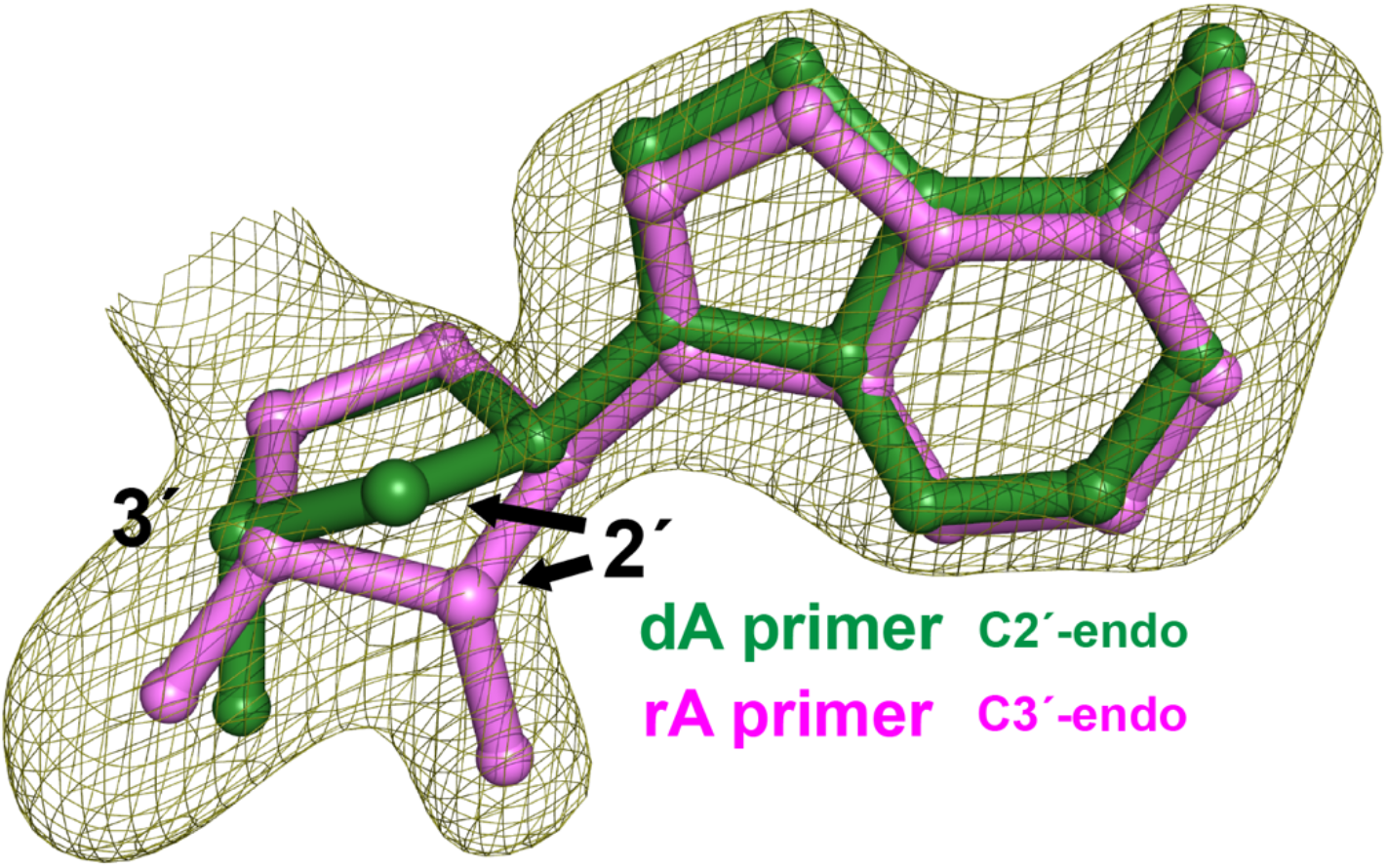
Structural comparison between primer terminus of dN versus rN extension. The F_o_-F_c_ omit map for the pink rA primer terminus was contoured at 3.5 σ.

**Fig. 6:**
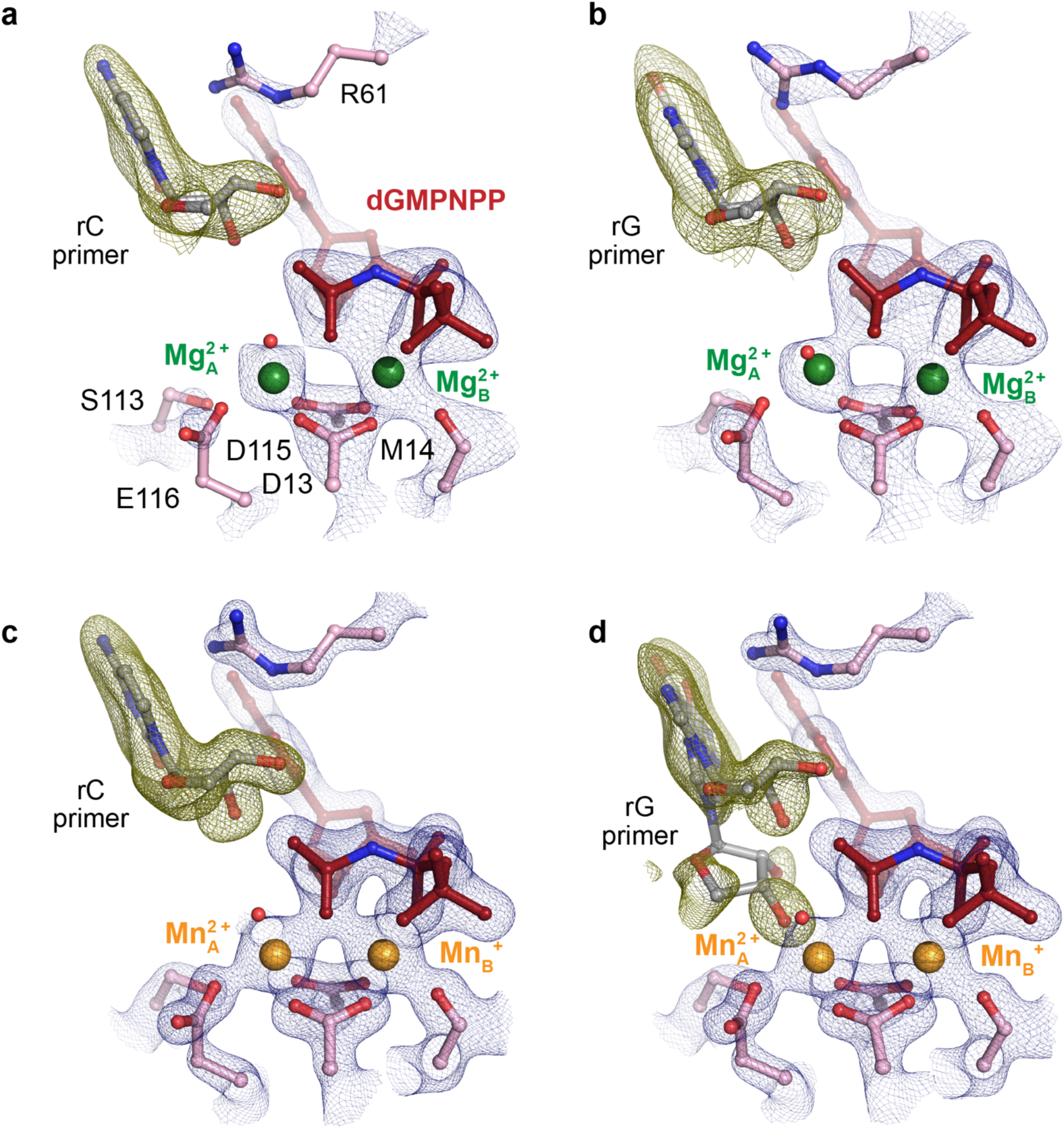
Structural comparison between reactant states of incorrect nucleotide incorporation during rC and rG extension. **a, b**, Structure of rC extension (**a**) and rG extension (**b)** misincorporation with Mg^2+^ by WT Pol η. **c, d**, Structure of rC extension (**c**) and rG extension (**d)** misincorporation with Mn^2+^ by Pol η. The 2F_o_-F_c_ map for everything including Me^2+^_A_ and Me^2+^ _B_, dGMPNPP, and catalytic residues and S113 (blue) was contoured at 2 σ. The F_o_-F_c_ omit map for the primer terminus (green) was contoured at 3.8 σ in **a** and **c** and 2.0 σ in **b**and **d**.

We suspect that the primer end is in dynamic equilibrium between the up and down conformations as observed in previous *in crystallo* studies (5,52). The occupancy might be too low for the down conformation of rC and rG to be refined at the current resolution. Because Mn^2+^ has been shown to improve primer alignment, we captured the same rC and rG-ended Pol η mismatch structures with Mn^2+^. The rG primed structure with Mn^2+^ showed increased presence of the primer terminus in the down conformation (from 0% to 30%) (**Fig. 6c and Supplementary Fig. 1**). The sugar pucker of the down conformation was also in C3’endo configuration. In contrast, for the structures with rC at the primer terminus, the primer remained in the up conformation, consistent with the kinetic studies that dC/rC-primed DNA exhibited the highest fidelity (**Fig. 6d and Supplementary Fig. 1**).

### *In crystallo rN extension* of Pol η

Previous studies have shown that nonhydrolyzable nucleotide analogs such as dNMPNPPs may impede primer alignment (5). Time-resolved crystallography allows the visualization of transient events that occur during biological reactions (3,4,48,52,55–60). Thus, we visualized the misincorporation process of deoxyribose guanine triphosphate (dGTP) across dT in the presence of Mn^2+^ from a single uridine primed DNA substrate by Pol η with a diffusion-based time-resolved crystallography system. In the ground state, 30% of the rU primer termini existed in the aligned down conformation and 3.6 Å away from the target α-phosphate (**Fig. 7a**). dGTP formed a wobble base-pair with the template dT. Similar to dT extension (5), 70% of R61 in the ground state was already flipped away from the triphosphate motif of the incoming nucleotide to stabilize the wobble dG-dT base pair.

**Fig. 7:**
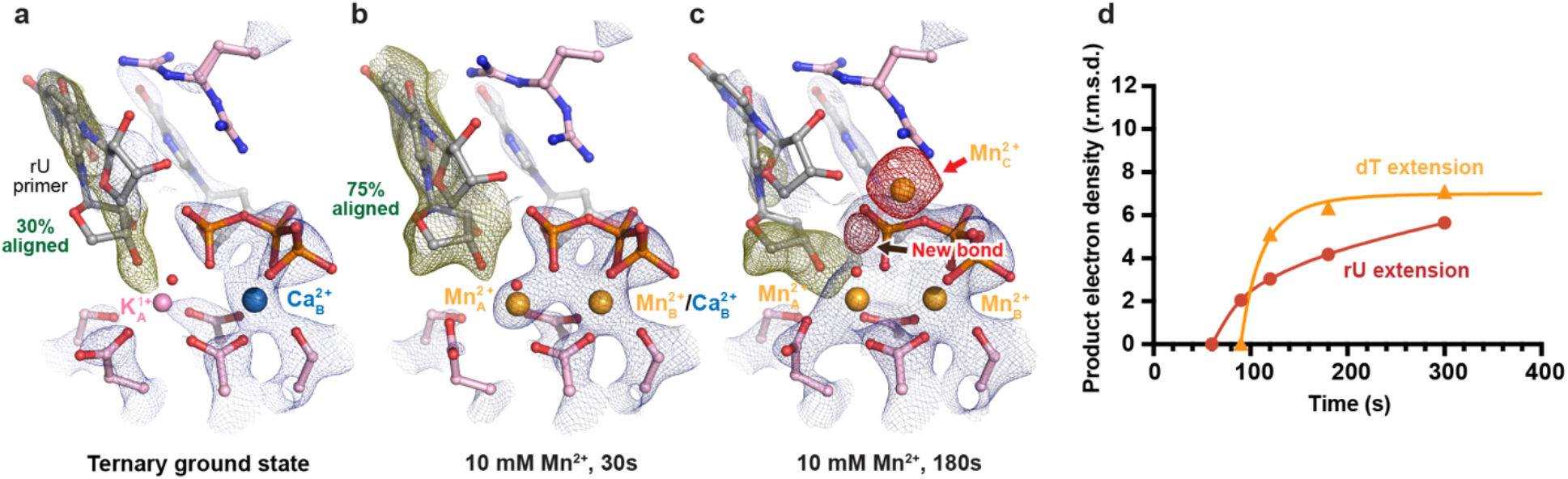
*In crystallo* visualization of WT Pol η rU extension misincorporation with Mn^2+^. **a-c,**Structures of WT Pol η during *in crystallo* catalysis after 10 mM Mn^2+^ soaking for 30s (**b**) and 180s (**c**). The 2F_o_-F_c_ map for the primer terminus up conformation (blue) was contoured at 1.2 σ. The 2F_o_-F_c_ map for the Me^2+^_A_, Me^2+^_B_, dGTP, and catalytic residues (blue) was contoured at 2.0 in **a**, 2.5 in **b**, and 2.0 σ in **c**. The F_o_-F_c_ omit map for the down conformation of the primer terminus (green) was contoured at 3 σ in **a-c**. The F_o_-F_c_ omit map for all the Mn^2+^_C_, and newly formed bond (red) was contoured at 3.5 σ. **d**, Timescale of reaction product of Pol η rU extension misincorporation *in crystallo* with Mn^2+^.

After 30s soaking in 10 mM Mn^2+^, 70% of the Me^2+^_A_ was already saturated with Mn^2+^, and 75% of the U primer had moved down in the aligned conformation, residing 3.6 Å away from the target α-phosphate (**Fig. 7b**). In comparison, after 30s soaking with a dT-ended primer, 65% of the Me^2+^_A_ was saturated with Mn^2+^ and 40% of the thymine primer was aligned for misincorporation. The Me^2+^_A_ during rU extension was assigned with 70% occupancy. Despite the improvement in alignment, the 3’-OH of rU resided 2.6 Å away from the Me^2+^_A_, 0.5 Å further compared to that during correct incorporation. After 180s of soaking, clear electron density for newly formed bond was visible and assigned to 50% between the rU 3’-OH and target α-phosphate (**Fig. 7c, d**). Unlike during dT extension in which the Me^2+^_C_ appeared before product formation, the Me^2+^_C_ here appeared simultaneously with product formation.

## Discussion

Many efforts have investigated the kinetic and structural mechanism surrounding polymerase fidelity (61–70). It was proposed that the exonuclease proofreading and the finger domains conformational changes (61,62,71,72) play significant roles in substrate discrimination. However, polymerases that lack proofreading exonuclease and finger domain conformational changes still incorporate correct bases versus incorrect bases more efficiently than what bases-pairing energies can provide (2,63,66,73–75). More recently, studies of Pol β and Pol η have shown that primer alignment contributes to the intrinsic polymerase fidelity and is perturbed during misincorporation (5,52). Here, we show that during rN extension, fidelity drops over 10-fold (**Fig. 1b**). Systematic structural studies confirmed that the drop in fidelity is likely due to improved primer alignment in the presence of incorrect incoming nucleotide (**Fig. 4,6,7**). Pol η fidelity is sequence dependent, with dA/dT at the primer end showing lower fidelity and dC with higher fidelity (53). The wobble base pair during rN extension looked identical to the previous reported dN extension mismatch structures, suggesting similar pi-stacking interactions (53). Consistent with the trend of fidelity, the rA and rU primer ends are better aligned compared to rC (**Fig. 4, 6**). These multiple level of correlation of primer end alignment and fidelity highlighted the critical role of primer alignment in polymerase fidelity control.

During DNA synthesis, the primer terminus overcomes a C2’-endo to C3’-endo barrier (3) to avoid steric clashes with the non-bridging oxygen of the incoming nucleotide. An extra 2’-OH on the sugar motif as in rN may affect sugar pucker conformation. Our structures show that rN primer termini are more readily in the C3’-endo form in the aligned conformation before product formation (**Fig. 5**). In addition, this suggests weaker barriers for product formation compared to DNA extension. The combination of improved primer alignment and weaker C2’-endo to C3’-endo sugar pucker conversion might explain the changes in fidelity. Many nucleoside analog drugs have modifications on the 3’ and 2’ carbon of the sugar ring and inhibit DNA synthesis at the extension step. Cytarabine, which is similar to dC but has a 2’-OH in the β direction, exists as a C2’-endo at the primer terminus even during correct nucleotide incorporation (76). The altered sugar ring conformation might increase barriers in C2’-endo to C3’-endo conversion and thus inhibit polymerase extension (76,77). Nucleoside analog drugs with altered sugar motifs such as entecavir and galidesivir may potentially exert their inhibitory effect through similar mechanisms.

Our studies on rN extension suggested the 2’-OH significantly affects polymerase fidelity. As Me^2+^_A_-mediated primer alignment is a required step in DNA synthesis, we hypothesize that the observed elevated incorporation error might be a universal property for all polymerases. However, polymerases from different families may evolve specific structural features to discriminate the 2’-OH at the primer end. Although Pol η and X-family polymerases can extend past rN primers with high efficiency, Pol ε efficiency decreases over 3300 fold during rN extension (36). Further biochemical and structural studies of different polymerases may be needed to clarify the effect of the 2’-OH group on catalysis, fidelity, and primer extension. The reduced efficiency of Pol ε in extending rN primer might indicate involvement of translesion polymerases in extending rN primer (5,52). In addition, the MMR pathway has been suggested to participate in rN removal but how MMR proteins recognize embedded rNs is not clear. It is possible that misincorporation during rN extension serves as a marker for exonuclease proofreading and MMR repair of rNs.

## Methods

### Protein expression and purification

Wild-type human polymerase η (Pol η) (residues 1–432) was cloned into a modified pET28p vector with a N-terminal 6-histidine tag and a PreScission Protease cleavage site as described (47). For protein expression, this Pol η plasmid was transformed into BL21 DE3 *E. coli* cells. When the optical density of the *E. coli* cells reached 0.8, isopropyl β-D-1-thiogalactopyranoside (IPTG) was added to a final concentration of 1 μM IPTG. After 20 hrs. (16 °C) of induction, the cell paste was collected *via* centrifugation and resuspended in a buffer that contained 20 mM Tris (pH 7.5), 1 M NaCl, 20 mM imidazole, and 5 mM ß-mercaptoethanol (BME). After sonification, Pol η was loaded onto a HisTrap HP column (GE Healthcare), which was pre-equilibrated with a buffer that contained 20 mM Tris (pH 7.5), 1 M NaCl, 20 mM imidazole, and 5 mM BME. The column was washed with 300 mL of buffer to remove non-specific bound proteins and was eluted with buffer that contained 20 mM Tris (pH 7.5), 1 M NaCl, 300 mM imidazole, and 3 mM dithiothreitol (DTT). The eluted Pol η was incubated with PreScission Protease to cleave the N-terminal 6-histidine-tag. Afterwards, Pol η was buffer-exchanged and desalted to 20 mM 2-(N-morpholino)ethanesulfonic acid (MES) (pH 6.0), 250 mM KCl, 10% glycerol, 0.1 mM ethylenediaminetetraacetic acid (EDTA), and 3 mM DTT and was loaded onto a MonoS 10/100 column (GE Healthcare). The protein was eluted with an increasing salt (KCl) gradient. Finally, Pol η was cleaned with a Superdex 200 10/300 GL column (GE Healthcare) with a buffer that contained 20 mM Tris (pH 7.5), 450 mM KCl, and 3 mM DTT.

### DNA synthesis assay

The nucleotide incorporation activity was assayed by the following: The reaction mixture contained 1.3-180 nM Pol η (WT or S113A), 5 μM DNA, 0-400 μM dNTP (either dATP or dGTP), 100 mM KCl, 50 mM Tris (pH 7.5), 5 mM MgCl_2_, 3 mM DTT, 0.1 mg/mL bovine serum albumin, and 4% glycerol. The incorporation assays were executed using DNA template and 5’-fluorescein-labelled primer listed in Table 1. Reactions were conducted at 37 °C for 5 min and were stopped by adding formamide quench buffer to the final concentrations of 40% formamide, 50 mM EDTA (pH 8.0), 0.1 mg/ml xylene cyanol, and 0.1 mg/ml bromophenol. After heating to 97 °C for 5 min and immediately placing on ice, reaction products were resolved on 22.5% polyacrylamide urea gels. The gels were visualized by a Sapphire Biomolecular Imager and quantified using the built-in software. Quantification of K_cat_, K_M_, V_Max_ and fitting and graphic representation were executed by Graph Prism. Source data of urea gels are provided as a Source Data file.

**Table 1.**
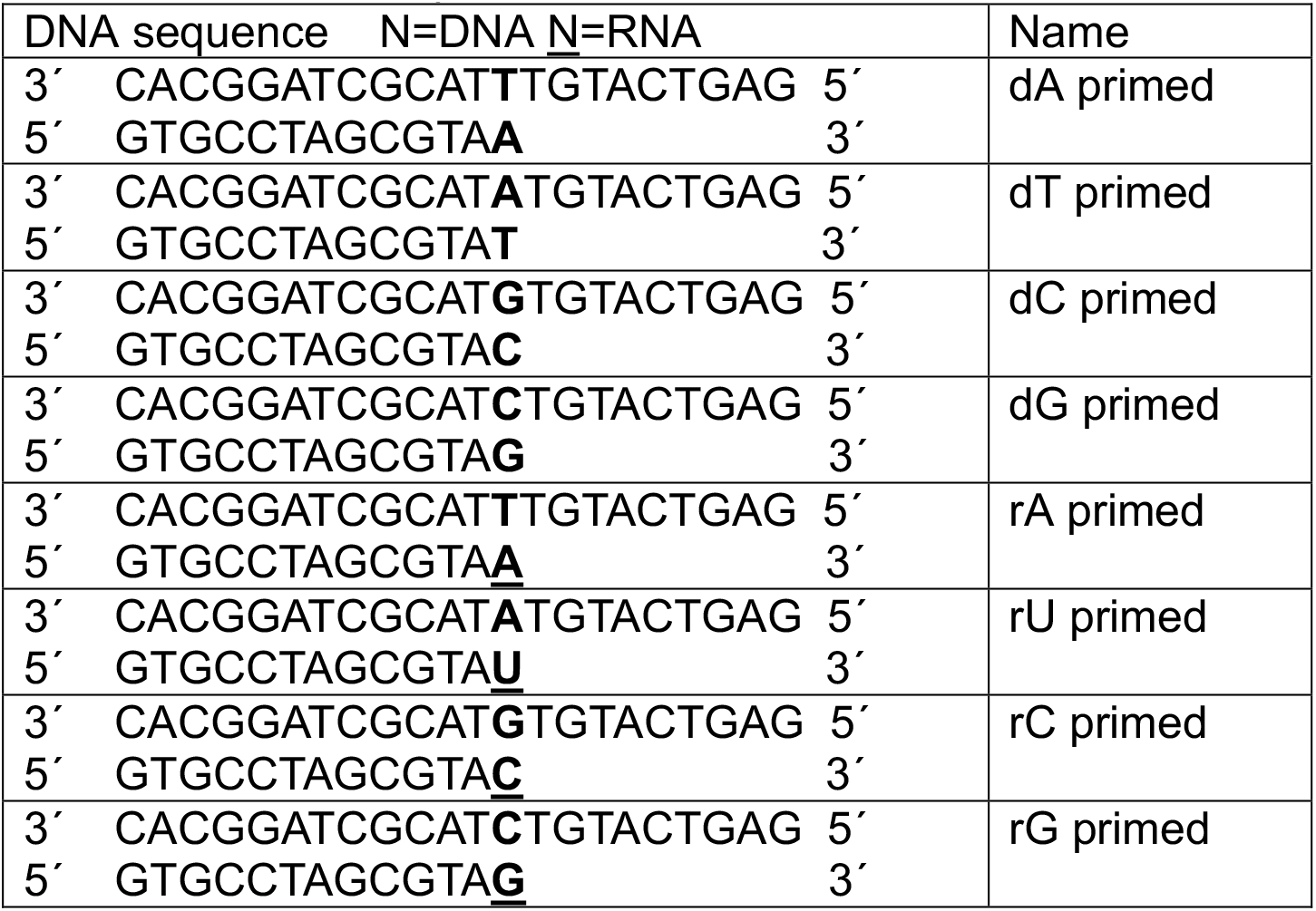
Kinetic Assay DNA sequences

### Crystallization

Pol η was concentrated to 300 μM in buffer that contained 20 mM Tris (pH 7.5), 0.45 M KCl, and 3 mM DTT. Then DNA, dGTP or dGMPNPP, and Ca^2+^ and low salt buffer (20 mM Tris (pH 7.5), and 3 mM DTT) were added to this polymerase solution at the molar ratio of 1 : 1.2 : 1 : 1 for Pol η, DNA, dGTP or dGMPNPP, and Ca^2+^, bringing Pol η’s concentration to 100 μM. Then after this solution was kept on ice for 10 min, more dGTP or dGMPNPP was added to a final concentration of 0.5 mM. DNA template and primer used for crystallization are listed in Table 2. All crystals were obtained using the hanging-drop vapour-diffusion method against a reservoir solution containing 0.1 M MES (pH 6.0) and 9-15% (w/v) PEG2K-MME at room temperature within 4 days.

**Table 2.**
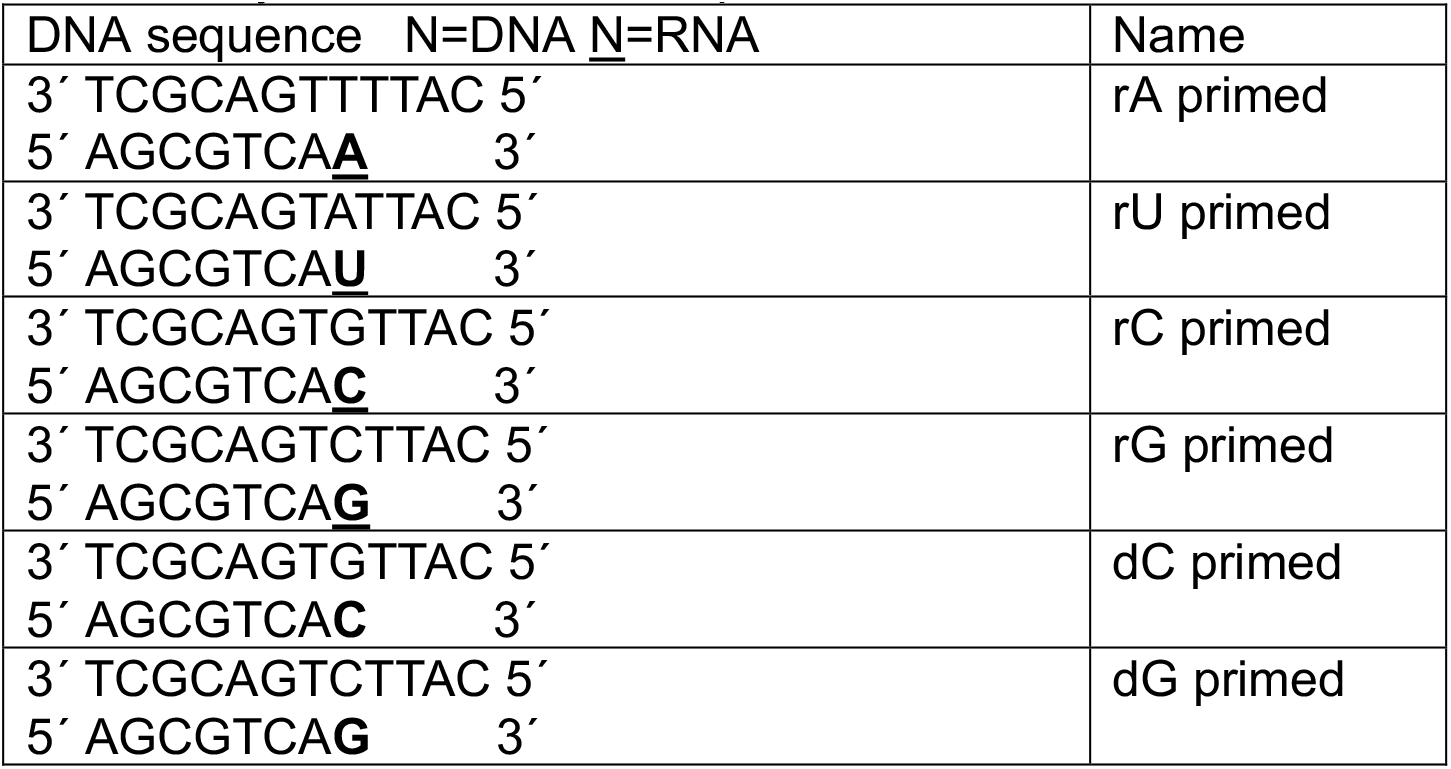
Crystallization DNA sequences

| DNA sequence | N=DNA | N=RNA | Name |
| --- | --- | --- | --- |
| 3' TCGCAGTTTTAC 5' |  |  | rA primed |
| 5' AGCGTCAA 3' |  |  |  |
| 3' TCGCAGTATTAC 5' |  |  | rU primed |
| 5' AGCGTCAU 3' |  |  |  |

|  |  |
| --- | --- |
| 3' TCGCAGTGTTAC 5'<br>5' AGCGTCAC 3' | rC primed |
| 3' TCGCAGTCTTAC 5'<br>5' AGCGTCAG 3' | rG primed |
| 3' TCGCAGTGTTAC 5'<br>5' AGCGTCAC 3' | dC primed |
| 3' TCGCAGTCTTAC 5'<br>5' AGCGTCAG 3' | dG primed |

### Chemical reaction *in crystallo*

The crystals were first transferred and incubated in a pre-reaction buffer containing 0.1M MES (pH 7.0, titrated by KOH), 100 μM dGTP, and 20% (w/v) PEG2K-MME for 30 min. The chemical reaction was initiated by transferring the crystals into a reaction buffer containing 0.1 M MES (pH 7.0), 20% (w/v) PEG2K-MME, and 10 mM MnCl_2_. After incubation for a desired time period, the crystals were quickly dipped in a cryo-solution supplemented with 20% (w/v) glycerol and flash-cooled in liquid nitrogen.

### Data collection and Refinement

Diffraction data were collected at 100 K on LS-CAT beam lines 21-D-D, 21-ID-F, and 21-ID-G at the Advanced Photon Source (Argonne National Laboratory). Data were indexed in space group P61, scaled and reduced using XDS (78). Isomorphous Pol η structures with Mg^2+^ PDB: was used as initial models for refinement using PHENIX (79) and COOT (80). Initial occupancies were assigned for the substrate, reaction product, PP_i_, Me^2+^_A_, Me^2+^_B_, and Me^2+^_C_, for the ternary ground state, following the previous protocol (4). After there were no significant F_o_-F_c_ peaks and each atom’s B value had roughly similar values to its ligand, we assigned occupancies for the same regions for the timepoints in between. Source data of the electron densities in r.m.s. density are provided as a Source Data file. Each structure was refined to the highest resolution data collected, which ranged between 1.75-2.2 Å. Software applications used in this project were compiled and configured by SBGrid (81). Source data of data collection and refinement statistics are summarized in **supplementary Table 1a-c**. All structural figures were drawn using PyMOL (http://www.pymol.org).

## Data availability

The coordinates, density maps, and structure factors for all the structures have been deposited in Protein Data Bank (PDB) under accession codes: 8E85, 8E86, 8E87, 8E88, 8E89, 8E8A, 8E8B, 8E8C, 8E8D, 8E8E, 8E8F, 8E8G, 8E8H, 8E8J, and 8E8K.

## Acknowledgements

Our sincere appreciation to the members of the Gao lab, and Drs. Phillips, Nikonowicz, and Lu who serve on CC’s thesis committee. We thank the APS LS-CAT beam technicians and research scientists Drs. Anderson, Wawrzak, Kondrashkina, and Focia. We thank Jamie Smith, Chuxuan Li, Joshua Chang for critical reading of the manuscript. This research used resources of the Advanced Photon Source, a U.S. Department of Energy (DOE) Office of Science User Facility operated for the DOE Office of Science by Argonne National Laboratory under Contract No. DE-AC02-06CH11357. Use of the LS-CAT Sector 21 was supported by the Michigan Economic Development Corporation and the Michigan Technology Tri-Corridor (Grant 085P1000817).

## Author contribution

YG conceived the project. CC carried out all time-resolved crystallography experiments, data collection and processing. CC and SE carried out protein purification, and CC and GZ executed protein crystallization. CC, CLL, SE, GZ, and AL performed the biochemical assays. CC and YG wrote the manuscript.

## Funding and additional information

This work is supported by CPRIT (RR190046) and the Welch Foundation (C-2033-20200401) to YG, and a predoctoral fellowship from the Houston Area Molecular Biophysics Program (NIH Grant No. T32 GM008280, Program Director Dr. Theodore Wensel) to CC.

## Supporting Information

**Fig. S1:**
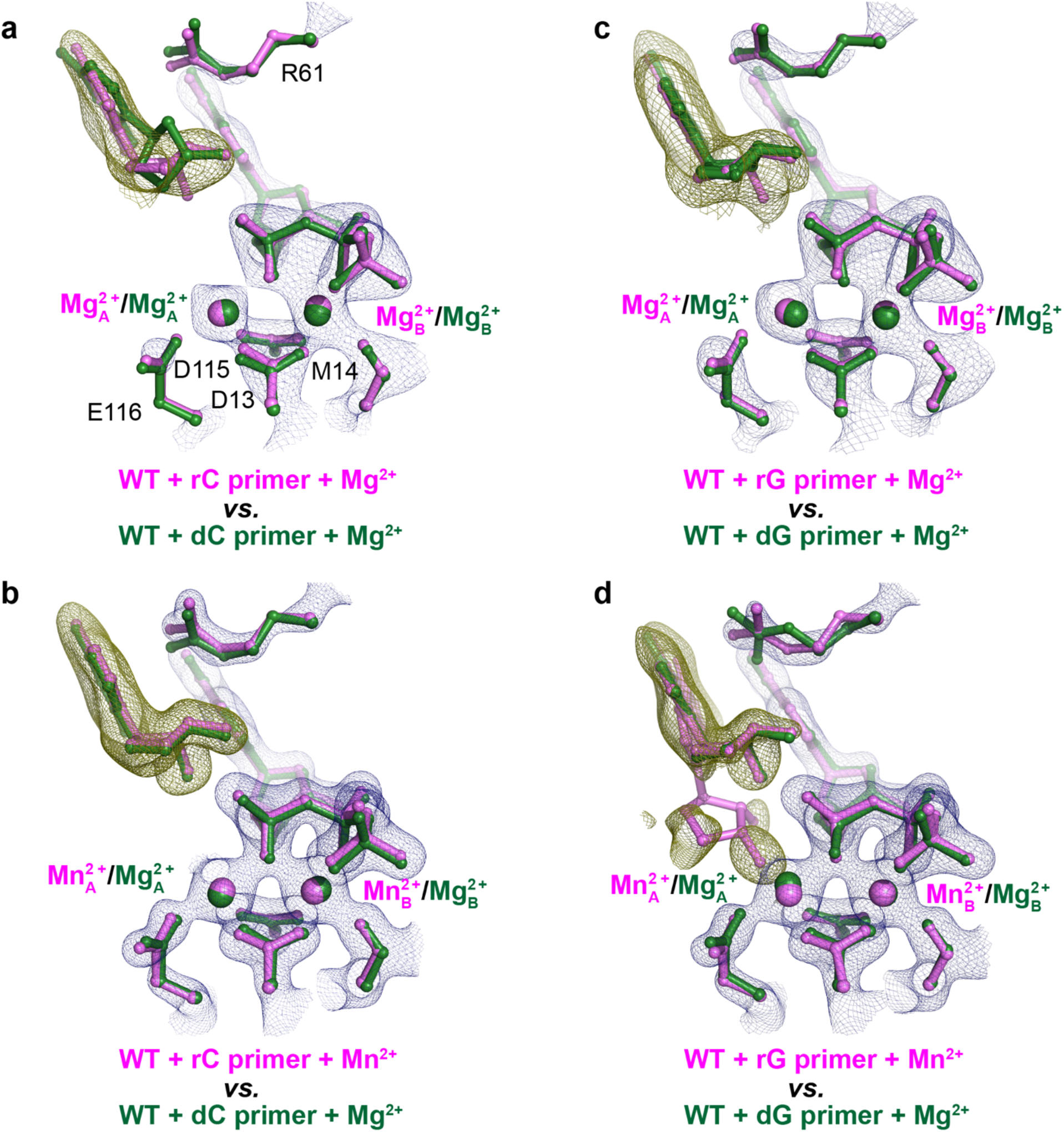
Structural comparison between reactant states of incorrect nucleotide incorporation during rC and rG extension. **a, b**, Structural overlay of dC (green) (PDB ID **4J9L**) or rC (pink) extension structures with dGMPNPP:dT base-pair. **c, d**, Structural overlay of dG (green) (PDB ID **4J9N**) or rG (pink) extension structures with dGMPNPP:dT base-pair All electron density maps apply to the molecule colored in pink. The 2F_o_-F_c_ map for everything including Me^2+^_A_ and Me^2+^_B_, dGMPNPP, and catalytic residues and S113 (blue) was contoured at 2.5 σ in **a, b** and 2 σ in **c, d**. The F_o_-F_c_ omit map for the primer terminus (green) was contoured at 3.8 σ in **a, b** and 2.0 σ in **c, d**.

**Table S1:**
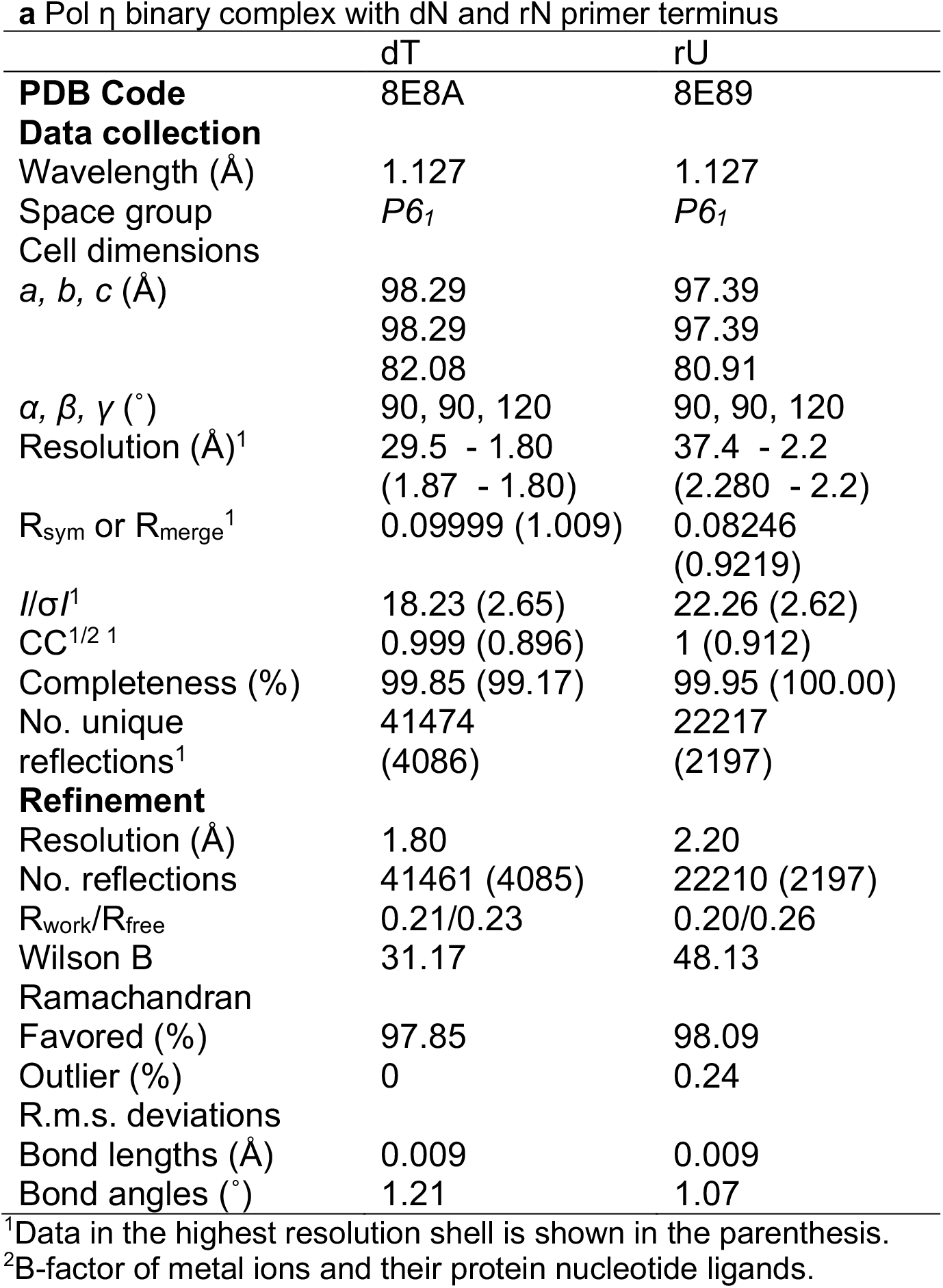
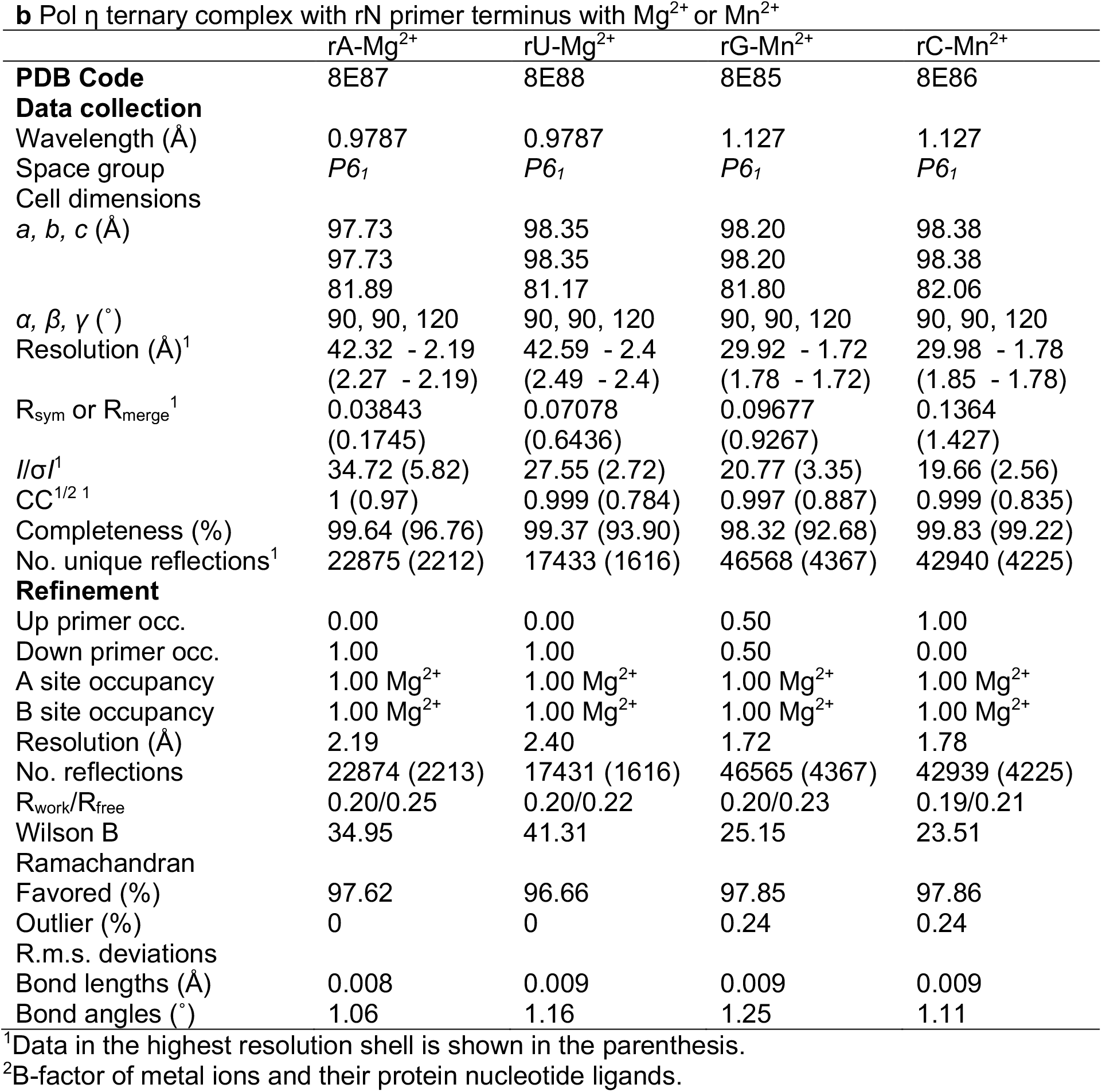
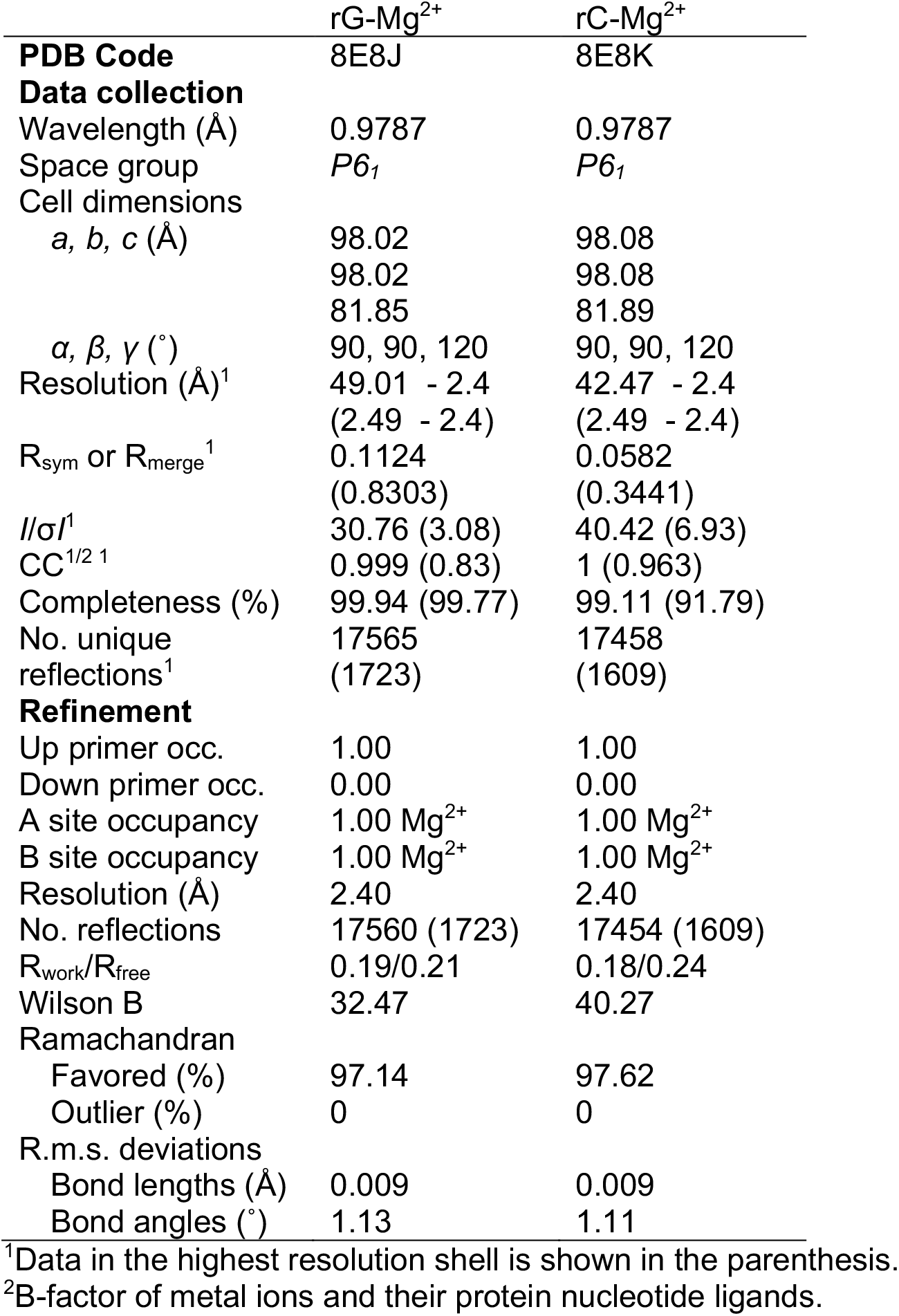
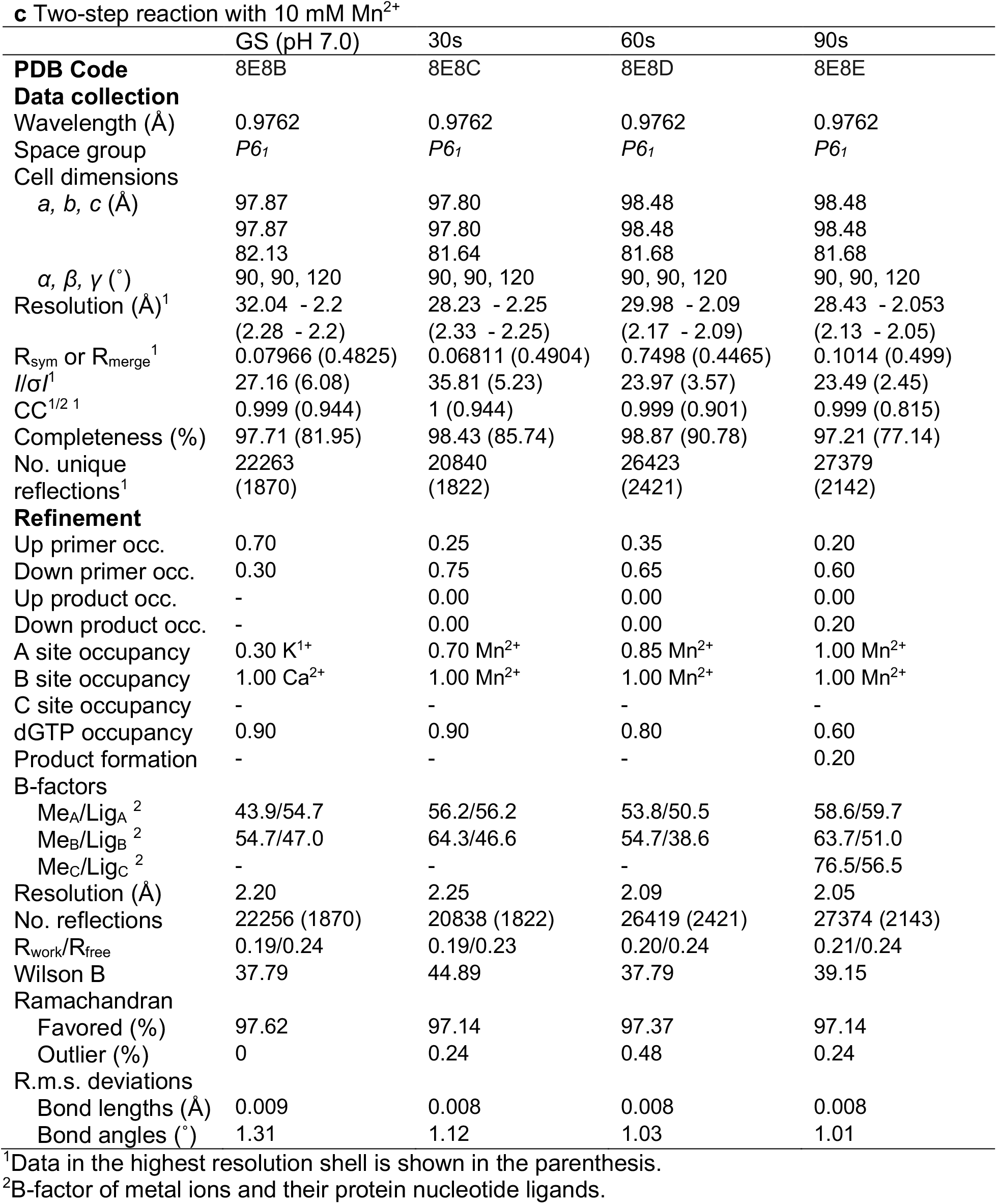
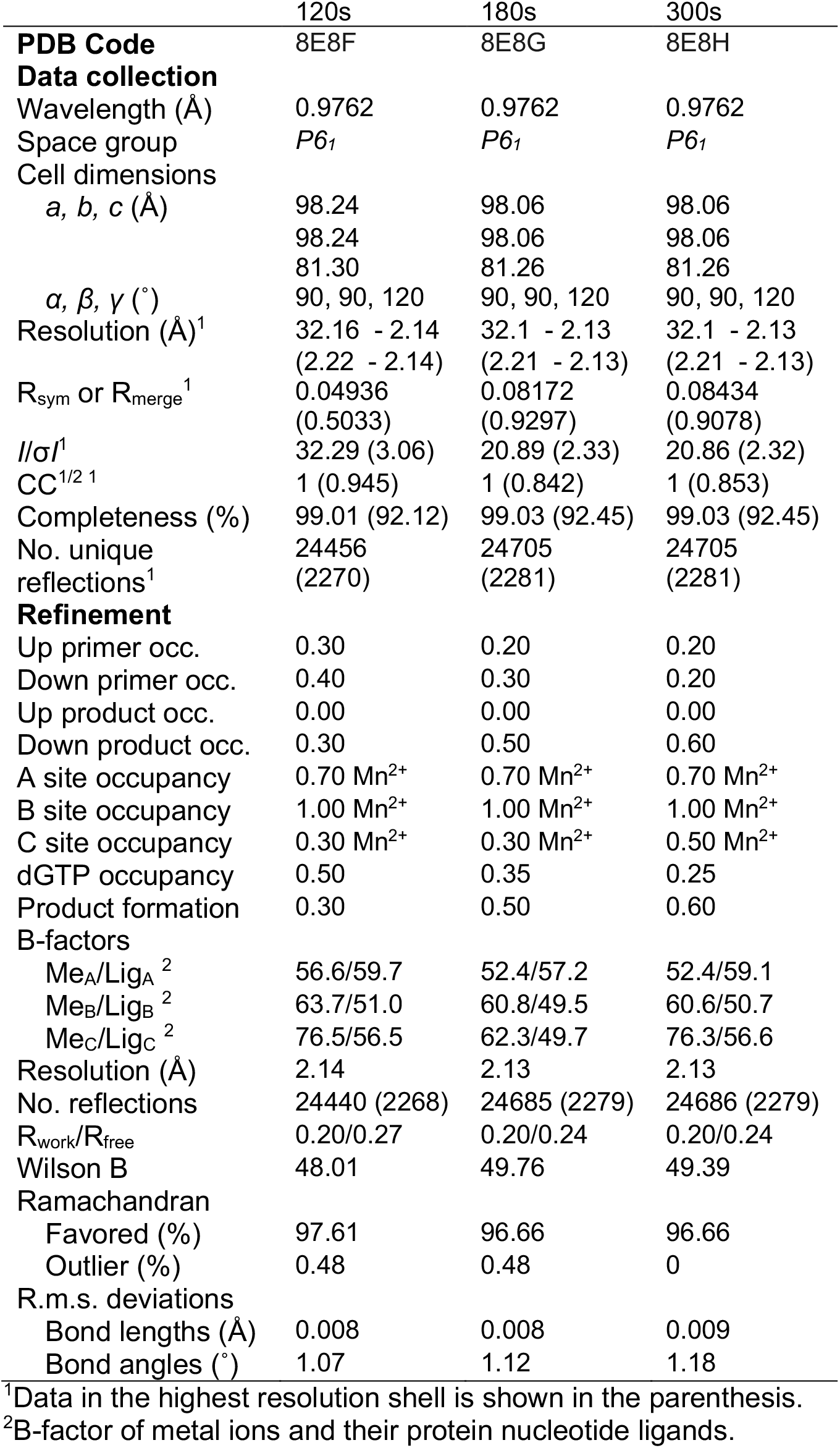
Crystal Diffraction and refinement data.

